# Different assembly and binning strategies recover contrasting genome catalogues and community profiles

**DOI:** 10.64898/2026.11.13.766504

**Authors:** Raphael Eisenhofer, Antton Alberdi, Ostaizka Aizpurua

## Abstract

In genome-resolved metagenomics, different assembly and binning strategies can be employed to reconstruct microbial genomes and community profiles. We used metagenomic data from a longitudinal experiment, in which mice were exposed to a series of environmental stressors, to compare four approaches: single-coverage, multi-coverage, co-assembly, and multi-split. We evaluated their effectiveness in reconstructing microbial genomes, determining their relative abundances, and analysing functional profiles while accounting for computational efficiency. Our findings indicate significant differences in the quantity and quality of recovered genomes across strategies, which in turn influenced community profiles and research outcomes related to microbiota resilience to environmental pressures and functional differences between exposure to contrasting temperatures. We conclude that the choice of assembly and binning strategies is crucial and needs to be thoroughly assessed considering available resources, study design, and research aims, as the strategy employed for data processing can significantly affect the results and conclusions of metagenomic studies.

**Importance:** Metagenomic sequencing allows researchers to reconstruct the genomes of the microbes living in an environment and to measure how abundant each one is. Before this can be done, the sequencing data must be assembled and grouped using one of several possible strategies, and the choice is often driven by habit or computational convenience. Using faecal samples from mice exposed to heat and cold, we show that ten strategies yield markedly different genome collections, community profiles, and even statistical conclusions about how gut microbiomes respond to environmental stress, while differing several-fold in computational cost. No single strategy performed best on all criteria. Our results offer quantitative guidance for choosing data-processing strategies that fit a study’s design, aims, and resources, and they warn that conclusions from metagenomic studies can depend on this often overlooked decision.

## Introduction

Genome-resolved metagenomics (GRM) has emerged as a common approach to characterise and analyse host-associated and environmental microbial communities (1, 2). The core of GRM involves bioinformatic reconstruction and annotation of bacterial and archaeal genomes from metagenomic mixtures (the so-called metagenome-assembled genomes or MAGs) (3), followed by quantification of their relative abundances across samples (4). The main strength of this approach lies in its ability to directly infer functionality from the gene contents encoded within the bacterial genomes under scrutiny (5), rather than relying on indirect inference, as in the case of 16S rRNA gene amplicon sequencing (6, 7).

Two critical steps of the GRM workflow are metagenomic assembly and binning (8). In the assembly step, overlapping reads are merged into longer contiguous sequences called contigs (9). Each contig may correspond to a portion of the genome of one or more microorganisms present in the sample. The subsequent binning step groups contigs into bins based on similarities in their sequence composition, read mapping coverage, and other features (10). The quality of these bins is then evaluated following multiple assessment criteria (11), with bins above a certain quality threshold being considered MAGs (3). The collection of MAGs reconstructed from a set of metagenomic samples becomes the catalogue of microbial genomes in the studied environment (12, 13), which is used as a reference for downstream diversity, compositional, and functional analyses (14). Consequently, the quality and representativeness of a MAG catalogue exert significant influence over the reliability and relevance of the ensuing results (11, 15).

While the overall workflow for reconstructing microbial genomes from metagenomic mixtures is relatively well-established (1, 8), researchers face critical decisions throughout the process. Regarding metagenomic assemblies, a key decision is whether to run individual assemblies per sample or to pool multiple samples into co-assemblies (16). Co-assemblies can improve the capture of low-abundance genomes (17), yet at the expense of collapsing strain diversity (18, 19). For binning, single- or multi-sample coverage strategies can be used, with multi-sample coverage providing higher-quality MAGs at the expense of computational resources (20). The performance of different metagenomic assemblers, binning algorithms, and bin refinement strategies has undergone extensive assessment using ground-truth benchmarks (10, 21–24). However, the extent to which the aforementioned decisions impact study results and, consequently, the conclusions drawn remains largely unexplored.

Here, we scrutinised the effect of multiple key decisions on GRM outputs and their effect on research conclusions based on metagenomic data generated in a longitudinal mice experiment (Fig. 1a). By combining four different approaches to assemble and bin a set of experimental samples (Fig. 1b) with three experimental design-specific sample grouping choices (Fig. 1c) (hereafter referred to as processing strategies, or simply, strategies; Table 1), we quantified the diversity and quality of reconstructed microbial genomes, measured the consistency of the recovered microbial communities, and critically assessed the impact of the alternative approximations on the generation and interpretation of biologically relevant results. In doing so, we aimed to raise awareness about the sensitivity of GRM approaches towards study design considerations, while equipping researchers with quantitative evidence to guide decision-making.

**Table 1.** Detailed description of each processing strategy.

| Processing strategy | Assemblies | Mapping | Binning |
|---|---|---|---|
| **single_coverage** | ***6 cages x 5 animals x 5 time-points***<br>One independent assembly per sample, for a total of 150. | ***(1 assembly) x 6 cages x 5 animals x 5 time-points***<br>Reads from each sample mapped only to its corresponding individual assembly. | Each assembly binned independently using the coverage information from its corresponding sample only. |
| **coassembly_animal** | ***6 cages x 5 animals***<br>30 co-assemblies generated by pooling reads from each mouse across all time points. | ***(5 time-points) x 30 co-assemblies***<br>Reads from each sample mapped individually to the corresponding animal-specific co-assembly. | Each animal-specific co-assembly binned using coverage information from all samples/time points collected from that animal. |
| **coassembly_timepoint_all** | ***5 time-points***<br>5 co-assemblies generated by pooling reads from all mice sampled at each time point. | ***(30 animals) x 5 co-assemblies***<br>Reads from each sample mapped individually to the corresponding time-point co-assembly. | Each per-time-point co-assembly binned using coverage information from all individual samples collected at that time-point. |
| **coassembly_timepoint_cage** | ***6 cages x 5 time-points***<br>30 co-assemblies generated by pooling reads per cage at time point. | ***(5 animals) x 30 co-assemblies***<br>Reads from each sample mapped individually to the corresponding cage x time-point co-assembly. | Each co-assembly binned using coverage information from all individual samples belonging to the corresponding cage x time-point group. |
| **multicoverage_animal** | ***6 cages x 5 animals x 5 time-points***<br>One independent assembly per sample, for a total of 150. | ***(5 time-points x 5 assemblies) x 30 animals***<br>Reads from all samples from each mouse mapped separately to each individual assembly from that mouse. | Each individual assembly binned using multiple coverage profiles obtained from all samples belonging to the same mouse. |
| **multicoverage_timepoint_all** |  | ***(30 animals x 30 assemblies) x 5 time-points***<br>Reads from all samples collected at the same time point were mapped separately to each individual assembly generated at that time point. | Each individual assembly binned using multiple coverage profiles obtained from all samples collected at the corresponding time point. |
| **multicoverage_timepoint_cage** |  | ***(5 animals x 5 assemblies) x 6 cages x 5 time-points***<br>Reads from all samples belonging to the same cage at the same time point were mapped separately to each individual assembly in that cage × time-point group. | Each individual assembly binned using the multiple coverage profiles from all samples belonging to the corresponding cage x time-point group. |
| **multisplit_animal** |  | ***(5 time-points x 1 concatenated assembly) x 30 animals***<br>All assemblies of an animal concatenated into one and reads from every time point mapped against it. | Each concatenated assembly per animal binned using multiple coverage profiles from all samples belonging to the same mouse. |
| **multisplit_timepoint_all** |  | ***(30 animals x 1 concatenated assembly) x 5 time-points***<br>All assemblies at a given time point concatenated into one and reads from every animal mapped against it. | Each concatenated assembly per time-point binned using multiple coverage profiles from all samples collected at the corresponding time point. |
| **multisplit_timepoint_cage** |  | ***(5 animals x 1 concatenated assembly) x 6 cages x 5 time-points***<br>All assemblies at a given time-point and cage concatenated into one and reads from the corresponding time-point and cage mapped against it. | Each concatenated assembly per cage and time-point binned using the multiple coverage profiles from all samples belonging to the corresponding cage x time-point group. |

**Figure 1.**
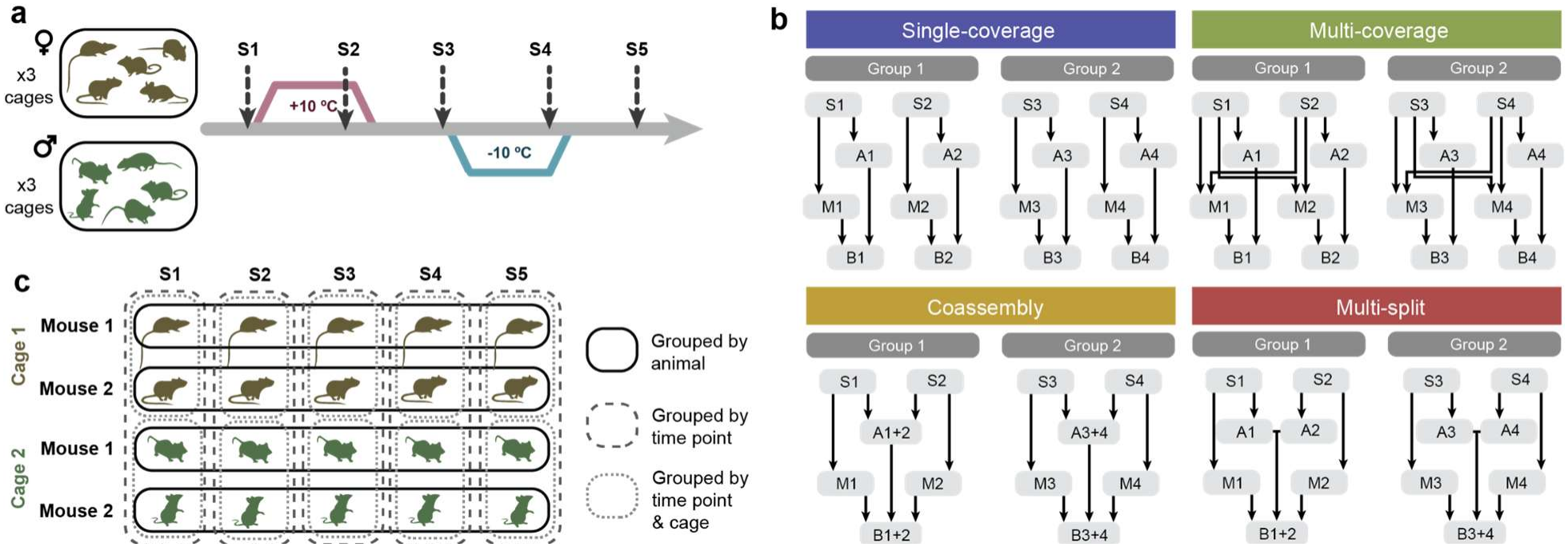
Experimental design and data processing strategies. **a)** Design of the laboratory experiment from which the samples were sourced. A total of 30 mice (15 females and 15 males) were housed in six unisex cages containing five mice each. Animals were first exposed to hot and then to cold temperatures, with faecal samples being collected at five time points before, during and after each environmental stressor. **b)** Overview of the four bioinformatic approaches compared in this study. Only two groups and two samples per group are shown for visualisation purposes. S: sample, A: assembly, M: mapping, B: binning. **c)** The three groupings that were compared in the study. Only two groups and two samples per group are shown for visualisation purposes.

## Results

Shotgun metagenome data were analysed using four bioinformatic approaches: single-coverage (individual assembly, mapping, and binning), multi-coverage (individual assembly and binning with multi-mapping), co-assembly (several samples are pooled for assembly and binning with individual mapping), and multi-split (concatenated individual assemblies, with reads mapped to the concatenated assembly and co-binned) (Fig. 1b). In addition, three groupings were used for data processing: (i) by animal (merging data from multiple time points), (ii) by time point (merging data from multiple individuals), and (iii) by time point and cage (merging data from each time point and cage) (Fig. 1c). By integrating different bioinformatic approaches and groupings, we assessed the impact of ten data-processing strategies (Table 1) on the generation and interpretation of biologically relevant results through quantification of the diversity and quality of reconstructed microbial genomes and evaluation of the consistency of the recovered microbial communities (Fig. 2a).

**Figure 2.**
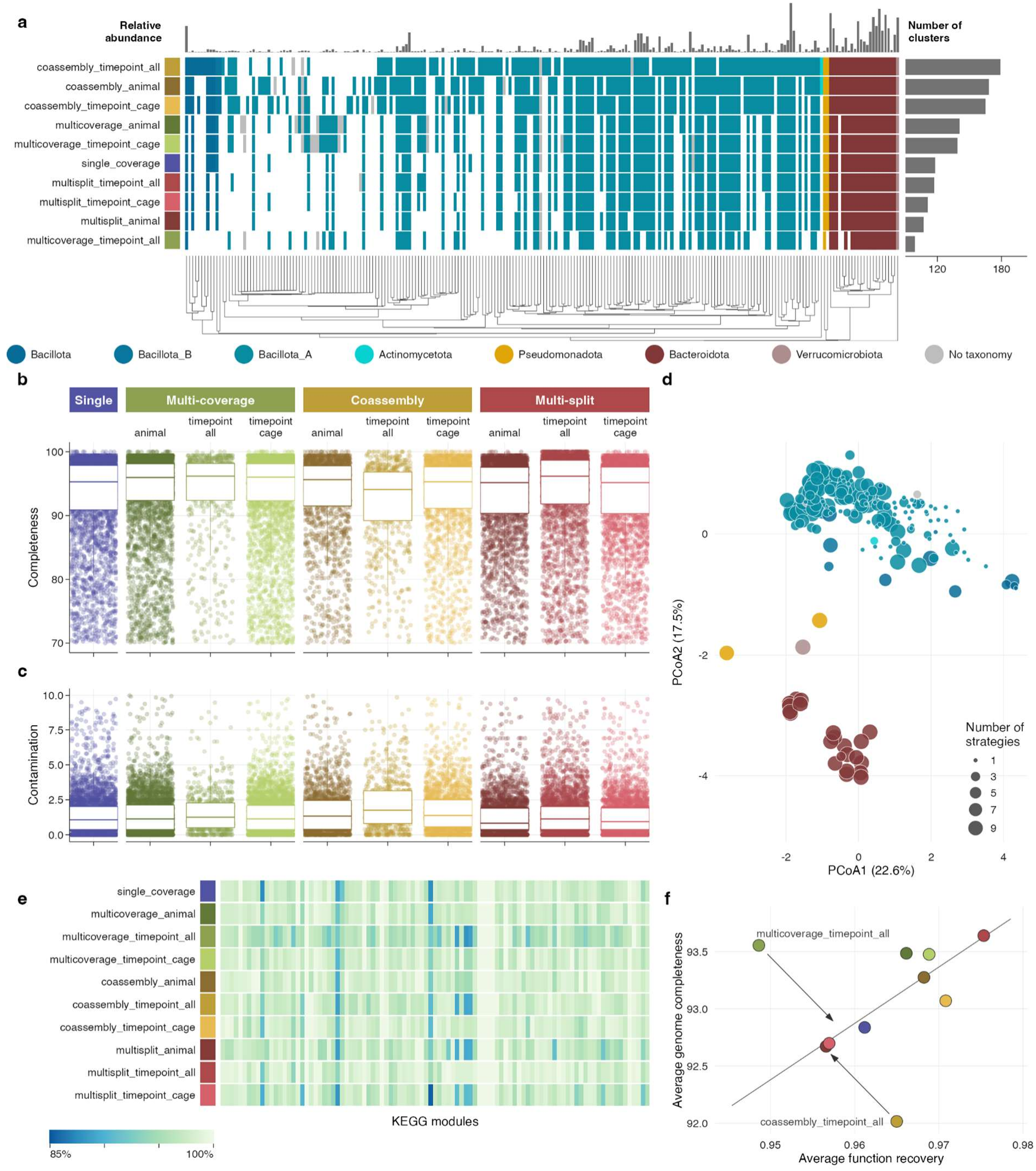
Quantitative and qualitative attributes of bacterial genome clusters recovered through different processing strategies. **a)** Overview of genome clusters recovered by each processing strategy. The vertical axis is sorted by the number of recovered clusters, while the horizontal axis is organised by microbial phylogeny. **b)** Average genome completeness estimations per processing strategy as reported by CheckM2. **c)** Average genome contamination estimations per processing strategy as reported by CheckM2. **d)** Ordination of genome clusters according to their functional attributes. Colours indicate different phyla, and dot sizes represent the number of processing strategies in which each genome cluster was recovered. **e)** Recovery of functional attributes by each processing strategy in terms of the percentage of module fullness compared to the maximum respective value obtained by any of the strategies. **f)** Relationship between the average function recovery and average genome completeness, showing the positive correlation between both attributes and highlighting the outlier strategies.

### Contrasting recovery of bacterial genomes

The ten data processing strategies yielded 25,983 redundant MAGs (CheckM2 completeness > 70%, contamination < 10%), which were grouped into 235 clusters (Fig. 2a). These clusters represent closely related genomes (> 98% average nucleotide identity) that belong to seven bacterial phyla, with Bacillota_A and Bacteroidota being the most broadly represented. None of the strategies recovered MAGs from all clusters, though MAGs from 77 clusters (32.8%) were retrieved by all strategies. The coassembly_timepoint_all strategy, which relied on five large assemblies (30 samples/assembly), yielded the highest number of clusters, confirming the importance of assembly size for recovering diversity (16). The next strategies with the highest number of clusters were the other two co-assembly approaches, which were based on a larger number of smaller assemblies (5 samples/assembly), underscoring the overall success of co-assembly approaches in recovering MAGs. In contrast, multi-split strategies, together with multicoverage_timepoint_all, recovered the lowest number of clusters. It is worth noting that the low performance of the multicoverage**_**timepoint_all strategy was likely due to only a subset of the samples being run through this approach due to large computational requirements (750 rather than the full 4,500 mappings; see Methods).

The likelihood of representation was significantly correlated with the relative abundance of the taxon (Spearman ρ = 0.88, p < 0.001). Highly abundant genomes were captured by all strategies, whereas rare genomes were captured by only a few or a single strategy (Supplementary Fig. S1). For instance, a single low-abundance Actinomycetota genome was retrieved by only two of the co-assembly strategies, while most of the highly abundant Bacteroidota were recovered by all strategies. Beyond assembly size, the higher performance of coassembly_timepoint_all might also be attributed to the presence of low-abundance taxa, some of them concentrated at a given time-point (e.g., hot exposure), which were only recovered when assembling data generated from each time point together (Supplementary Fig. S2). The lower cluster recovery of the various multi-split and single-coverage approaches relative to co-assembly could be due to the reliance on individual assemblies, which, in this study, probably suffered from insufficient sequencing depth (3.6±0.7 GB/sample or 12.3±2.5 million read pairs/sample) for adequate assembly coverage.

Quality metrics of recovered MAGs also differed across strategies (Fig. 2b-c). The average CheckM2 completeness value was 93.14±6.92%, while the average CheckM2 contamination was 1.48±1.47% (Supplementary Fig. S3). The processing strategy that yielded the largest number of clusters (coassembly_timepoint_all) had the lowest completeness and highest contamination values. In contrast, multisplit_timepoint_all, which yielded one of the lowest numbers of clusters, exhibited the highest completeness values. Discrepancies in the taxonomic classifications of contigs within bins were calculated using GUNC, with multi-split strategies yielding the highest percentage of bins passing GUNC (98-99%), compared to co-assembly approaches (95-97%; Supplementary Fig. S4).

The reconstructed genomes were clustered based on their KEGG metabolic module fullness attributes (Supplementary Fig. S5) into two main functional groups that aligned with phylogeny (Fig. 2d). Bacteroidota, Pseudomonadota and Verrucomicrobiota formed a dense group of functionally similar genomes, whose members were recovered by most strategies. In contrast, the highly diverse Bacillota_A formed a wider functional group along with Bacillota, Bacillota_B and Actinomycetota, with many clusters only recovered by one or a few strategies due to their lower abundances. Completeness of MAGs exerted a direct impact on the recovered functional attributes: function recovery was highest in multisplit_timepoint_all and lowest in multicoverage_timepoint_all (Fig. 2e). The recovery of functional traits was found to be positively correlated with MAG completeness in all strategies, in agreement with previous observations on the impact of genome completeness on functional inference (25). However, multicoverage_timepoint_all exhibited a functional recovery rate that was significantly lower than expected given its high completeness values, while coassembly_timepoint_all yielded the opposite result, recovering more functions than its lowest completeness would predict. This indicates that incomplete genomes impacted functional inference less than expected in the latter strategy (Fig. 2f).

### Contrasting recovery of community attributes

All but the multisplit_animal strategy reached top domain-adjusted mapping rates (DAMR) (26), indicating a nearly complete recovery of expected microbial reads by their respective genome catalogues. This observation likely results from the increased average completeness compensating for the lower numbers of MAGs in strategies that yielded fewer MAGs. Additionally, this effect was probably amplified by the limited read recruitment of rare genomes, which were only recovered by some strategies (Supplementary Fig. S6), compared to the extensive recruitment of abundant genomes, which were recovered by all strategies (Supplementary Fig. S1).

Despite similar DAMRs, the recovery of diverging numbers of MAGs exerted an impact on the quantitative attributes of microbial communities. We analysed alpha and beta diversity metrics within the Hill numbers framework to account for different components of diversity (27–29): only presence-absence (richness), presence-absence and evenness (neutral), and presence-absence, evenness and phylogenetic relationships among MAGs (phylogenetic). The effect of processing strategies was statistically significant in all three diversity metrics analysed (linear mixed-model LMM_ANOVA_; richness F[9, 893]=553.19, neutral F[9, 893]=331.64, phylogenetic F[9, 893]=74.50; p < 0.001), with co-assembly strategies recovering higher diversities, although differences tended to decrease as more components of diversity were considered (Fig. 3a). Despite the differences in alpha diversities, single_coverage yielded very similar microbial communities to co-assembly strategies in terms of compositional differences (Fig. 3b). In contrast, multi-split and multi-coverage strategies reconstructed considerably different community compositions, with dissimilarity patterns varying across diversity metrics.

**Figure 3.**
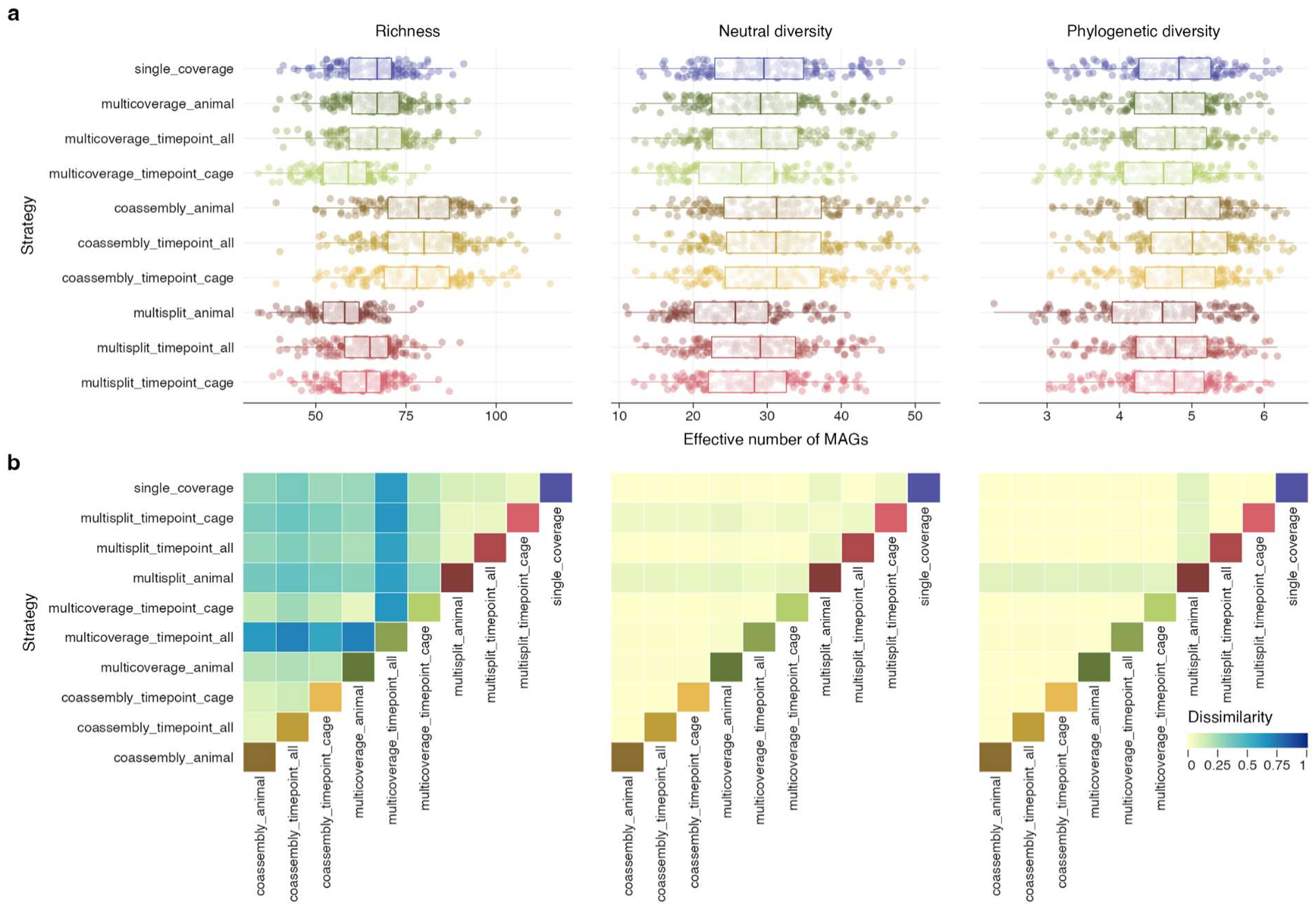
Impact of processing strategies on microbial community attributes. Diversity metrics account for different components of diversity: presence-absence only (Richness), presence-absence and evenness (Neutral diversity), and presence-absence, evenness and phylogenetic relationships among genomes (Phylogenetic diversity). **a)** Average alpha diversity values for the different metrics. **b) (**Dis)similarities between community ordinations constructed from each processing strategy according to the procrustes test.

### Impact on biological inference

We conducted two hypothesis-driven analyses to assess the impact of processing strategies on biological inference, underscoring the significance of these strategies in determining statistical support and shaping study conclusions.

The first analysis investigated whether the microbiota of female and male mice exhibited different levels of resilience to environmental stress. We measured the compositional turnover of microbiomes at three time points: before heat stress, between heat and cold stress, and after cold stress. Although females generally showed higher overall stability, the difference between sexes was statistically significant (uncorrected for multiple testing) in only six out of the ten processing strategies when considering richness, and in none when considering neutral or phylogenetic turnover (Fig. 4a). Additionally, overall turnover values varied between strategies, with co-assemblies revealing higher overall dissimilarities compared to other strategies.

**Figure 4.**
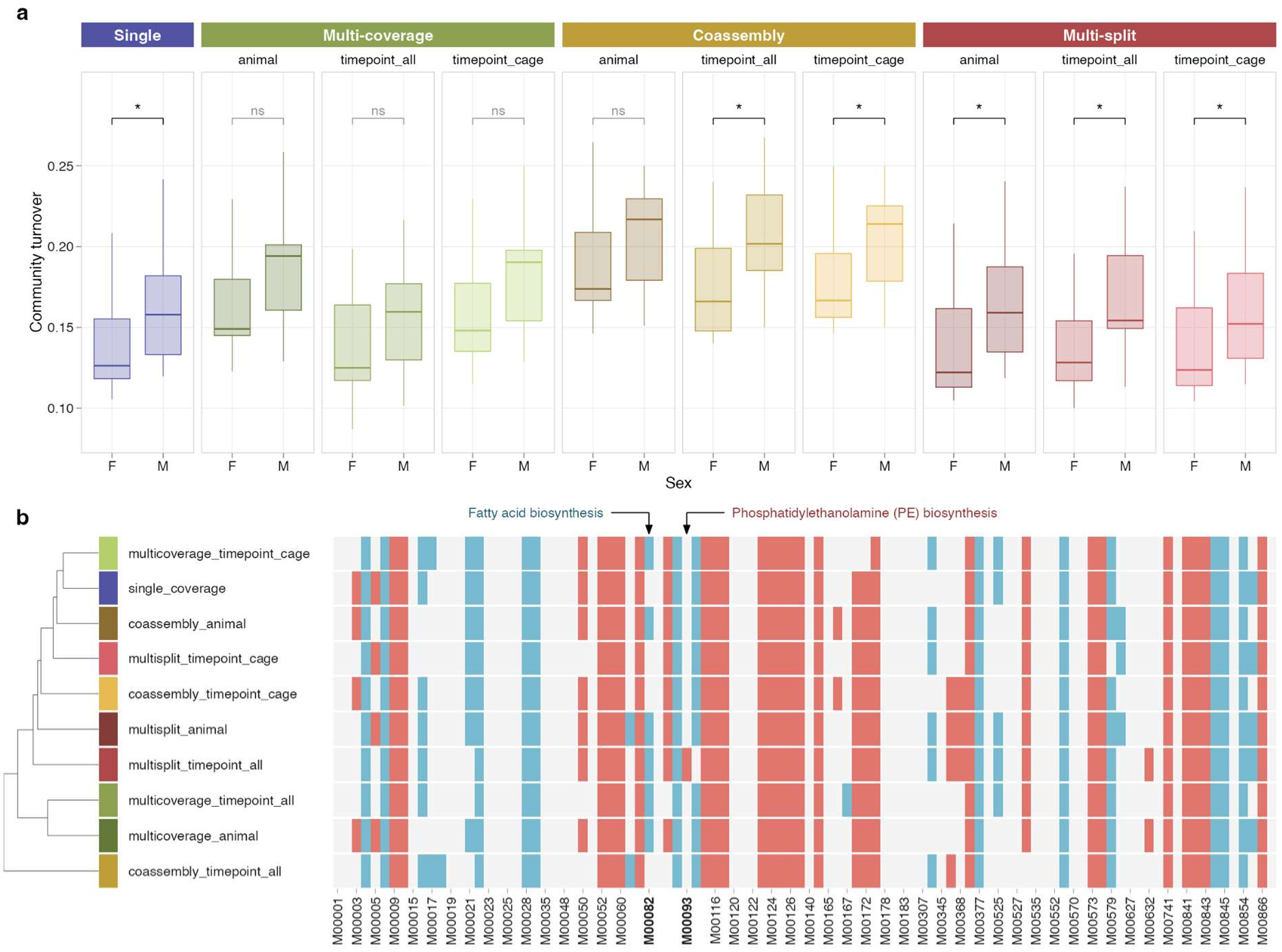
Impact of processing strategies on biological significance. **a)** Comparison of community turnover between female and male mice across various processing strategies. Asterisks (*) denote statistically significant differences according to the Kruskal-Wallis test (α=0.05), which varied among the processing strategies. **b)** Comparison of community-weighted average KEGG metabolic module fullness levels in microbial communities under cold and hot environmental conditions. Colours represent statistical significance of linear mixed models: blue tiles indicate significantly higher fullness levels under cold conditions, while red tiles indicate significantly higher fullness levels under hot conditions. Modules M00082 and M00093 are highlighted for illustrative purposes.

The second analysis compared the functional microbiota profiles of mice exposed to hot and cold temperatures. Specifically, we contrasted the community-weighted average fullness values of KEGG metabolic modules between microbial communities of each individual using linear mixed models, with animal identity as a random effect. While none of the strategies produced contradictory results, 25% of the modules exhibited differences across strategies (Fig. 4b). For example, M00082, which describes fatty acid biosynthesis, was significantly higher under cold conditions in six out of ten strategies. In contrast, M00093, which describes the biosynthesis of the membrane phospholipid phosphatidylethanolamine, remained invariant across all strategies except for multisplit_timepoint_all. The multisplit_animal strategy identified significant differences in 53% of the modules, whereas the multicoverage_timepoint_all strategy reported differences in only 44% of the modules. Overall, the coassembly_timepoint_all strategy yielded the most distinct results compared to the other strategies.

### Cost-effectiveness of data processing

Based on the above-presented results, one could conclude that combining multiple processing strategies (e.g. co-assembly with multi-coverage) is the best way to construct the most complete MAG catalogues and representative community profiles. However, data processing comes with a significant computational burden, requiring longer data processing times. To assess this, we benchmarked the computational resources needed to generate the data using the different strategies. We found that the multi-coverage strategies took on average 3.6 times longer than co-assemblies (2.3–4.3 times depending on the grouping strategy) to process the data (Supplementary Fig. S7a). This was largely driven by the increase in the number of mapping and bin refinement jobs. In an attempt to generalise our estimations, and based on the average computational requirements of the multiple bioinformatic steps, we also estimated the computational time needed to generate data using the different strategies under different study designs and sample sizes (Supplementary Fig. S7b). Single-coverage and co-assembly mostly increased linearly with an increasing number of samples and groups, whereas multi-coverage, and to a lesser extent multi-split, increased polynomially (quadratically in multi-coverage and approximately as n^1.6^ in multi-split). Note that these estimates are for samples with an average of 3.6 GB of data each, and as such, samples with more data (and higher genomic diversity) are expected to take longer.

## Discussion

Our scrutiny of assembly and binning strategies revealed significant differences in the resulting genome catalogues, their functional features, and their quantitative community profiles. In addition, we showed that these differences can lead to different study conclusions, and that the computational resources required for generating the data can vary drastically. Co-assembly approaches are computationally lightweight and recover large diversities, but at the expense of lower-quality genomes. In contrast, multi-coverage and multi-split strategies can reconstruct higher-quality genomes, but fail to recover the rarest genomes. In our study, single_coverage exhibited intermediate values in most analysed metrics. Our results might differ when applied to study systems with varying levels of microbiome complexity and sample structure. Nonetheless, our findings underscore the importance of data processing strategy choice while highlighting the impossibility of finding a one-size-fits-all solution. Therefore, we encourage researchers to carefully consider their metagenomic data processing strategy in light of their study design, research objectives, data volume, and computational resources.

## Materials and Methods

### Data processing

Shotgun raw metagenomic data archived within BioProject PRJEB77276 were used in this study. The dataset originates from an experiment conducted on outbred mice, in which 30 individuals (15 females and 15 males) were housed in six unisex cages, each containing five mice. Following an acclimation period, animals were systematically exposed to two environmental treatments: hot and cold stress. Each treatment lasted 10 days and was separated by a 4-day transitional phase. At each sampling point, individual mice were briefly isolated to ensure accurate sample collection independent of cage-level mixing.

To reduce differences in sequencing depth across cages, samples from mice with the highest sequencing yields were downsampled before downstream analysis. Downsampling was performed selectively to bring sequencing depths to approximately 3.5–4.5 GB per sample, with the aim of minimising differences in the average sequencing depth between animals and cages while retaining as much sequence information as possible. This procedure was used to reduce sequencing-depth imbalance as a potential source of bias in comparisons among processing strategies. All reads were subsampled to the desired base counts using BBMap’s reformat.sh (https://github.com/BioInfoTools/BBMap/tree/master). Primary bioinformatic processing was performed using the EHI Bioinformatics pipeline (30). Briefly, reads were quality-trimmed using fastp (31) before being mapped to the *Mus musculus* reference genome (GRCm39) using Bowtie2 (32). Host reads were filtered from BAM files using samtools (33). Non-host reads were then either assembled individually or co-assembled in groups (Fig. 1) using MEGAHIT (34). Binning of individual assemblies (single-coverage and multi-coverage) and co-assemblies was then performed using CONCOCT (35), MaxBin2 (36) and MetaBAT2 (37), before being refined using MetaWRAP’s bin_refinement module (1). Multi-split binning was performed using VAMB (38), as this is the only widely used binner that implements this approach. In consequence, the differences attributed to the multi-split approach are partially confounded with the choice of binning tool and cannot be fully disentangled in the present design.

Bins with CheckM2 completeness >70% and contamination <10% were retained as MAGs. MAGs were dereplicated at 98% ANI using dRep (18) with MASH (39) and FastANI (40), aiming to find a balance between strain-level resolution without the confounding effect of read cross-mapping across genomes. Dereplication was run separately for each strategy to generate strategy-specific catalogues, and jointly across all strategies to define the genome clusters compared among strategies. Non-host reads were mapped to the dereplicated MAG catalogue of each strategy using Bowtie2, with coverage statistics calculated using CoverM (41). Domain-adjusted mapping rates (DAMR) (26) were calculated for each sample and strategy as the percentage of reads mapped to the MAG catalogue divided by the percentage of bacterial and archaeal reads estimated with SingleM. We note that the multicoverage**_**timepoint_all strategy was only run with a subset of the data (5 mice * 5 time points * 30 assemblies = 750 mappings), to avoid the prohibitive need for 4,500 mappings the full dataset would require. MAGs were annotated taxonomically using GTDB-Tk v2.3.0 (42) with the GTDB r214 (43). Genome completeness and contamination (Fig. 2b–c) were estimated with CheckM2 (11), and chimerism was additionally assessed using GUNC (44). DRAM (5) was used to annotate MAG functions (45–52). Functional attributes were expressed as KEGG module fullness (the proportion of steps of each module present in a genome), taking for each genome cluster and strategy the maximum fullness among its MAGs. Genome clusters were ordinated through principal coordinates analysis (PCoA) based on Euclidean distances between their module fullness profiles. Function recovery was calculated for the clusters recovered by all strategies as the fullness of each module in a given strategy relative to the maximum value obtained by any strategy, averaged across modules. The relationship between average function recovery and average genome completeness across strategies was assessed through linear regression and Pearson correlation, and the two strategies with the largest residuals were highlighted.

### Data analysis

Diversity analyses were based on Hill numbers and were computed using the R package hilldiv2 v2.0.5 (53). For all alpha and beta diversity measurements, we calculated three metrics accounting for different components of diversity: presence-absence only (Richness), presence-absence and evenness (Neutral diversity), and presence-absence, evenness and phylogenetic relationships among genomes (Phylogenetic diversity). Alpha diversity differences between processing strategies were analysed through linear mixed modelling as implemented in lmerTest (54), using sample identity as a random effect. F-statistics were subsequently computed using the Satterthwaite method. Beta dissimilarity metrics were computed for all pairwise combinations in terms of Sørensen-type local overlap-complement using hilldiv2 (53). Subsequently, NMDS ordinations were conducted using metaMDS, and differences between ordinations were tested using protest (55).

For the first hypothesis-driven analysis, we computed community variation metrics across the three time points in which the animals were housed under optimal conditions (T1, T3 and T5). The turnover was measured in terms of Sørensen-type local overlap-complement, and mean values were compared between females and males using the Kruskal-Wallis H test (equivalent to the Wilcoxon rank-sum test for two groups), without correction for multiple testing and without accounting for the cage in which animals were housed (sex and cage are confounded in this design, as cages were unisex).

For the second hypothesis-driven analysis, we contrasted functional differences between the hot and cold exposure time points (T2 and T4) through linear mixed modelling as implemented in lmerTest (54), using the animal identity as a random effect. Community-weighted mean fullness was calculated for each sample as the sum of the module fullness values of all genome clusters weighted by their relative abundances. Only modules with a community-weighted mean fullness above 0.3 were tested, and p-values were Bonferroni-corrected across modules within each strategy.

Computational requirements were derived from the Snakemake benchmark files (wall-clock time per job) of the assembly, mapping, binning and bin refinement steps, and total processing time per strategy was calculated as the sum of job times. Times for alternative numbers of samples and groups were extrapolated from the mean per-job times, assuming linear scaling of assembly, binning and refinement with the number of samples, and quadratic scaling of mapping in multi-coverage strategies.

## Data, Metadata, and Code Availability

Raw sequencing data are archived at the ENA under BioProject PRJEB77276. Code, metadata, and data to reproduce these analyses are available at https://github.com/alberdilab/assembly_binning_strategies, with a snapshot archived in Zenodo with DOI 10.5281/zenodo.24074915.

## Acknowledgments

This study was funded by the Danish National Research Foundation grant DNRF143.

